# Improving emotion control in social anxiety by targeting rhythmic brain circuits

**DOI:** 10.1101/2023.09.01.555689

**Authors:** Sjoerd Meijer, Bob Bramson, Ivan Toni, Karin Roelofs

## Abstract

Social avoidance is a hallmark of social anxiety disorder. Difficulties in controlling avoidance behavior are the core maintaining factor of this impairing condition, hampering the efficacy of existing therapies. This preregistered study tested a physiologically-grounded non-invasive enhancement of control over social approach and avoidance behavior in socially anxious individuals. Their prefrontal and sensorimotor areas received dual-site phase-coupled electrical stimulation, to enhance endogenous inter-regional theta-gamma phase-amplitude coupling, a mechanism known to support emotion control in non-anxious individuals. We measured behavioral and fMRI-BOLD responses during in-phase, anti-phase, and sham stimulations, while participants performed a social approach-avoidance task, involving either automatic or controlled emotional actions. In-phase stimulation selectively enhanced control over approach-avoidance actions, and modulated neural responses in the same prefrontal region where stimulation-reactivity increased as a function of trait anxiety. These findings illustrate how human neurophysiological connectivity can be leveraged to improve control over social avoidance, opening the way for mechanistically grounded clinical interventions of persistent avoidance in anxiety disorders.

**SIGNIFICANCE STATEMENT:** Controlling automatic approach-avoid behavior is essential for human social interactions. This ability is impaired in social anxiety, but we know how neurotypical brains use endogenous rhythmic coupling for social emotion control. In a preregistered study, we used noninvasive electrical brain stimulation to enhance endogenous rhythmic coupling between prefrontal theta- and sensorimotor gamma-band rhythms, while highly socially anxious individuals solved an emotional control challenge. In-phase stimulation selectively enhanced emotional action control and modulated neural activity in a prefrontal cortex region where brain stimulation reactivity increased as a function of trait anxiety. These findings provide evidence for the generalizability and clinical potential of this interventional approach to social anxiety.

## INTRODUCTION

Anxiety disorders are marked by an inability to flexibly regulate prepotent approach and avoidance tendencies to adapt behavior to contextual demands. Persistent avoidance of perceived threats causes and maintains anxiety, preventing the extinction of those threats through exposure, thereby exacerbating the disorder (Lovibond et al., 2009; Pittig et al., 2015; Craske and Stein, 2016; Rattel et al., 2017). There is an urgent need for non-invasive interventions targeted at improving emotional action control in anxious individuals. Recent work has improved cognitive and behavioral functions in neurotypical individuals, boosting inter-regional neural communication through dual-site phase-amplitude-coupled transcranial alternating current stimulation (dual-site tACS) (Polanía et al., 2012; Violante et al., 2017; Preisig et al., 2021; Grover et al., 2022). This non-invasive neuromodulation technique, grounded on the physiology of directional inter-regional neuronal communication, can also enhance emotion control in non-anxious individuals (Bramson et al., 2020a). Here, we test the generalizability of this approach by applying dual-site tACS in a group of participants selected for high social anxiety, a trait known to be associated with impaired emotional action control (Heuer et al., 2007; Roelofs et al., 2009b; van Peer et al., 2009). This translational test is a critical step toward developing clinically-relevant interventions for modulating emotional behavioral control.

Patients suffering from social anxiety disorder have particular difficulties in overriding automatic avoidance tendencies in social situations. Previous research has identified the lateral prefrontal cortex (lPFC) as a critical brain region involved in selecting between competing approach-avoidance actions (Koch et al., 2018; Bramson et al., 2020b). This control is implemented through rhythmic modulation of excitability in downstream areas, including the sensorimotor cortex (SMC) and parietal cortex (Voytek et al., 2015; Bramson et al., 2018, 2020a; Weber et al., 2024). Specifically, successful control over social-emotional approach-avoidance actions has been linked to coupling between the phase of theta-band rhythms in lPFC and the amplitude of gamma-band activity in SMC (Voytek et al., 2015; Bramson et al., 2018; Weber et al., 2024).

Dual-site tACS offers a means to non-invasively modulate endogenous inter-regional phase-amplitude coupling, enhancing long-range synchronization between task-relevant neural circuits (Reinhart and Nguyen, 2019; Bramson et al., 2020a; Grover et al., 2022). Our work has shown that in-phase stimulation, designed to synchronize peaks of theta-band rhythms in lPFC with the amplitude of gamma-band neural activity in SMC, improves accuracy when non-anxious participants override automatic emotional action tendencies (Bramson et al., 2020a). Given the pivotal role of the lPFC in regulating approach-avoidance actions (Volman et al., 2011) and the importance of theta-gamma coupling between lPFC and SMC for successful control over social-emotional approach-avoidance actions (Bramson et al., 2018, 2020a), here we test whether dual-site tACS applied to lPFC-SMC can enhance emotional action control in highly socially anxious individuals. We applied in-phase and anti-phase dual-site tACS while forty-nine highly socially anxious individuals performed an approach-avoidance task, using concurrent fMRI to quantify prefrontal engagement during control over emotional action tendencies.

## MATERIALS AND METHODS

### Participants

Fifty-two highly socially anxious students (39 females) from Radboud University, Nijmegen, were recruited to participate in this pre-registered experiment (https://osf.io/j9s2z/). The local ethics committee approved the experiment (METC Arnhem-Nijmegen: CMO2014/288). Two participants were excluded due to malfunctioning of the stimulation equipment, and one participant was excluded for failing to comply with task instructions during one of the stimulation sessions. Participants were selected based on high self-reported social anxiety levels, as detailed below, and screened for eligibility criteria, including no history of mental illness other than anxiety disorders or current use of psychoactive medication. Additionally, all participants had normal or corrected-to-normal vision and underwent screening for contraindications to magnetic resonance imaging (MRI) and non-invasive transcranial electrical stimulation. The mean age of participants was 24.0 years (SD = ±4.3), with an age range of 19 to 40 years.

Pre-screening was conducted using the Liebowitz Social Anxiety Scale (LSAS). Participants who scored 30 or higher (on the combined fear/anxiety and avoidance sub-scales) were included, as this cutoff provides an optimal balance between sensitivity and specificity for identifying individuals who meet the criteria for social anxiety disorder (Mennin et al., 2002). This selection criterion resulted in a normal distribution of LSAS scores within our study sample, with a mean of 64.4, a standard deviation (SD) of ±18.8, and a range from 31 to 106 (**Fig. 1A**). The trait sub-scale of the State-Trait Anxiety Inventory (STAI Y-2) was also included to assess trait anxiety, serving as an inter-individual difference marker of tACS BOLD dose-response. This is because endogenous prefrontal activation in response to the emotional action control challenge varies as a function of trait anxiety (Bramson et al., 2023).

**Figure 1.**
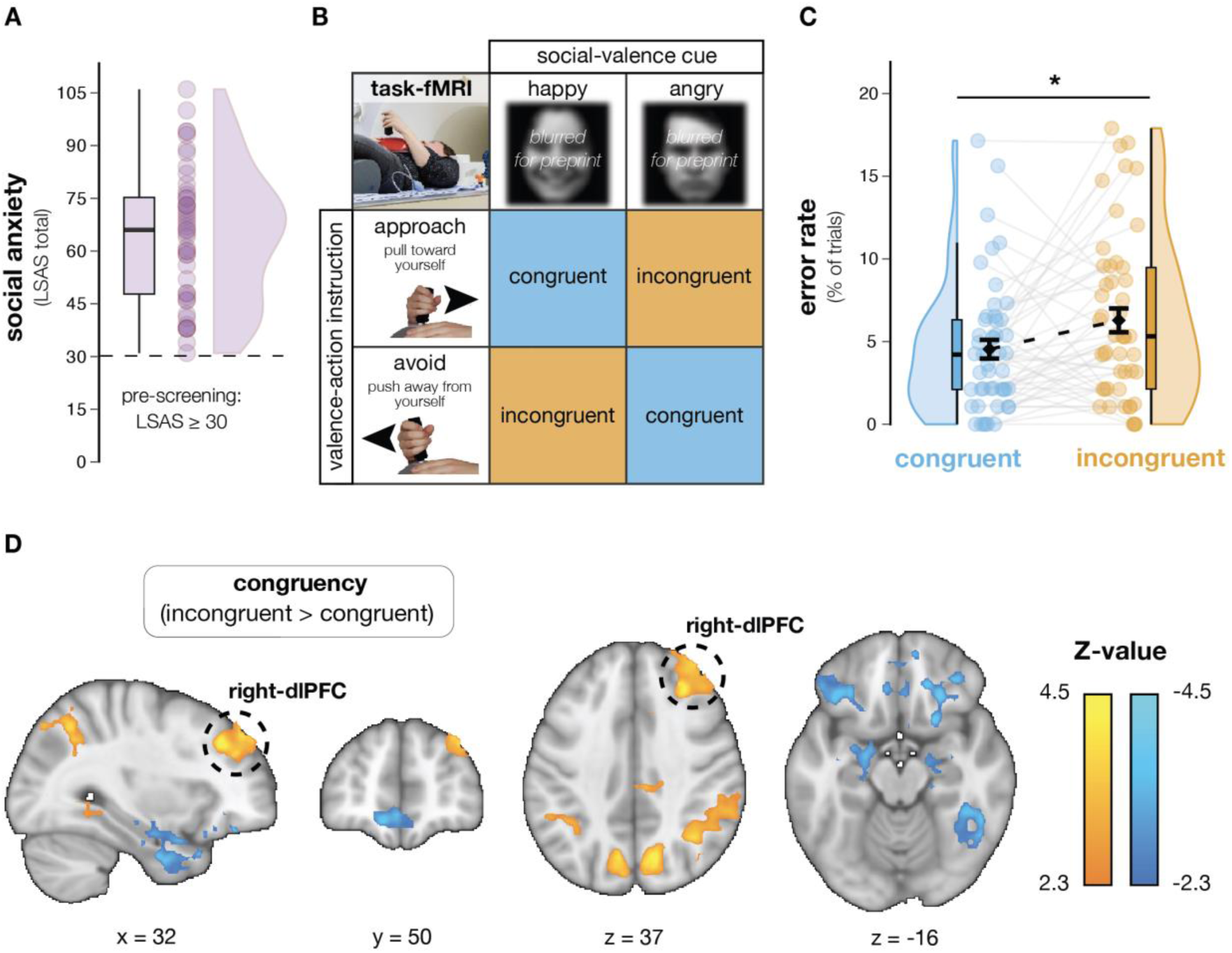
**Behavioral and BOLD fMRI effects related to control over emotional action tendencies. (A) Sample distribution of self-reported Liebowitz Social Anxiety Scale (LSAS) scores.** The black dashed line marks an LSAS score of 30, used to select socially anxious participants. This cut-off balances specificity and sensitivity in detecting heightened social anxiety (Mennin et al., 2002). **(B) Conceptual visualization of the approach-avoidance task performed during concurrent dual-site tACS and fMRI.** In affect-congruent conditions, participants were instructed to pull the joystick towards them in response to a happy face and push it away from them in response to an angry face. In contrast, the affect-incongruent condition requires participants to override automatic emotional action tendencies to avoid angry and approach happy faces. Therefore, in this condition, participants were instructed to pull the joystick towards them in response to an angry face and push it away in response to a happy face. **(C) Behavioral congruency-effects in the baseline sham stimulation condition of the task.** Participants made more errors when overriding their automatic emotional action tendencies in the affect-incongruent condition (orange circles) as compared to making affect-congruent responses (blue circles). Black-filled diamonds represent mean values, and standard errors of the mean are shown with black lines. **(D) Task-fMRI congruency-related effects across tACS stimulation conditions (incongruent > congruent; cluster-level inferences corrected for multiple comparisons).** Stronger BOLD signals were observed in prefrontal- and parietal areas when automatic emotional action tendencies needed to be overridden, while amygdala-hippocampal and medial-orbitofrontal areas showed weaker BOLD signals.

The current study in highly socially anxious individuals was powered to replicate the main finding of our previous investigation in healthy male students (Bramson et al., 2020a) that showed tACS phase- and dose-specific effects on emotional action control (https://osf.io/j9s2z/). This required a sample size that allowed us to detect the previously observed significant three-way interaction between Congruency (congruent, incongruent) * tACS-phase (in-phase, anti-phase) * tACS-dose (BOLD signal in lPFC during stimulation *vs*. sham): *η*_p_^2^ = 0.19 (Bramson et al., 2020a). The minimum sample size required for this study to detect an effect of comparable magnitude is 45 participants (*a priori* sensitivity power analysis of 2×2 RM-ANCOVA: α = .05, power = .80, *η_p_*^2^ = 0.19; G*Power v3.1(Faul et al., 2007)). Given that the current sample might be more heterogeneous due to mixed-sex and anxiety-related differences, we decided to increase our planned sample size to 50 participants.

### Procedure

The data collection procedure took place over three separate days. On the first day, participants completed the Liebowitz Social Anxiety Scale (LSAS) and State-Trait Anxiety Inventory-Trait (STAI-trait, Y-2) questionnaires. Following this, participants underwent a series of scanning procedures, including a structural T1-weighted scan, a diffusion-weighted imaging scan (DWI, as reported in (Bramson et al., 2020a, 2023)), and a magnetic spectroscopy scan (MRS, as reported in (Bramson et al., 2020a, 2023)).

On the second and third days of the experiment, neuronavigation was employed to position the electrodes over the sensorimotor cortex (SMC) and lateral prefrontal cortex (lPFC). Detailed information on electrode placement is provided in the following section. During both stimulation sessions, participants were positioned inside the MR scanner, where they first completed a 5-minute practice task. After the practice session, participants proceeded with the approach-avoidance task, which was conducted concurrently with dual-site tACS intervention and task-based functional MRI for approximately 35 minutes.

### Approach-avoidance task

Emotional action control was manipulated using a validated social approach-avoidance task in which cognitive control is associated with lPFC-SMC theta-gamma coupling (Bramson et al., 2018), and is sensitive to the phase-dependent effects of dual-site theta-gamma phase-amplitude tACS when controlling for inter-individual variation in prefrontal BOLD response to tACS (Bramson et al., 2020a). During functional MR scanning, participants lay inside the MR scanner with a joystick in their right hand resting on their lower abdomen. Participants were instructed to approach or avoid by moving the joystick toward or away from themselves, respectively. During the task, participants received written instructions on the screen (>10 s) at the start of each block of 12 trials (∼60 s) that stated, *“Pull the joystick toward yourself when you see a happy face and push away from yourself when you see an angry face”* (congruent condition), *“Push the joystick away from yourself when you see a happy face and pull toward yourself when you see an angry face”* (incongruent condition). Congruent and incongruent conditions alternated between blocks in a pseudo-random fashion. Between blocks, there was an inter-block interval of >20 s. Each trial started with a fixation cross (500 ms) followed by a face stimulus (100 ms) to which participants had to respond within 2 s. Joystick movements over 30% of the maximum movement range were considered valid responses. If a participant did not respond, they would receive on-screen feedback stating, *“You did not move the joystick far enough”*. Instructions and the inter-block interval lead to an ∼30 s wash-out period between stimulation conditions. Each participant had 288 trials on each of the two testing days, yielding 576 observations (trials) in total, equally distributed over congruency (congruent, incongruent) and stimulation (in-phase, anti-phase, sham) condition combinations (96 trials per bin).

### Dual-site tACS stimulation

Dual-site tACS was applied online during task performance inside the MRI scanner using two sets of concentric ring electrodes (inner disc: 25 mm ø; outer ring: 80 mm inner-ø, 100 mm outer-ø) (Saturnino et al., 2017). Current direction alternated (−1 mA to +1 mA) between each disc-ring electrode set to achieve a time-varying electrical field in theta-band (6 Hz) frequency over right-lPFC and gamma-band (75 Hz) tapered with a 6 Hz theta wave over left-SMC. The tACS-phase manipulation was achieved by phase-locking gamma-band power to peaks (in-phase) or troughs (anti-phase) of the theta-band signal. During sham blocks, there was an initial 10 s period of stimulation to match potential sensations related to the onset of stimulation, after which stimulation was terminated. Within a session, participants received all stimulation conditions (in-phase, anti-phase, sham) alternating in a pseudo-random fashion between stimulus blocks of 12 trials (∼60 s), interleaved with periods of no stimulation (instructions between blocks; > 30 s), and never repeating the same stimulation condition for two consecutive blocks.

We used neuronavigation (Localite TMS Navigator; RRID: <u>SCR_016126</u>) for individualized targeting of left-SMC (MNI [-28, -32, 64] (Bramson et al., 2018, 2020a)) and right-lPFC (MNI [26, 54, 0] (Neubert et al., 2014; Bramson et al., 2018, 2020a)) based on anatomical masks of regions of interest registered to individual T1-weighted scans acquired during the first session. After localization, electrodes were attached to the participant’s scalp using Ten20 conductive paste (MedCaT). Inside the MR scanner, stimulation was delivered using two Neuroconn DC-Stimulator Plus stimulators (neuroConn; impedance < 10 kOhm; RRID: <u>SCR_015520</u>). Stimulators were placed inside a magnetically shielded box designed with electronics that filtered out RF pulses of the MR scanner and combined with a BrainAmp ExG MR amplifier (www.brainproducts.com) that allowed continuous monitoring of stimulation output during the session.

### Modeling of stimulation currents

The electric field magnitude (mV/mm) at the cortical target sites was estimated using SimNIBS (version 4.0; (Thielscher et al., 2015)). The concentric ring electrode sets were modeled as the disc (25 mm ø) and outer ring electrode (80 mm inner-ø, 100 mm outer-ø) with the same center, using a 2-layer medium (2 mm silicone rubber on top of 2 mm saline gel). The electrode sets were positioned over lPFC and SMC using the template head mesh of SimNIBS. For all media, we used standard conductivities provided by SimNIBS. Direct currents of 1 mA were modeled to flow between the center disc electrode and the outer ring. The model results of the electric field magnitude at the cortical surface for both target sites are shown in **Fig. 2A**.

**Figure 2.**
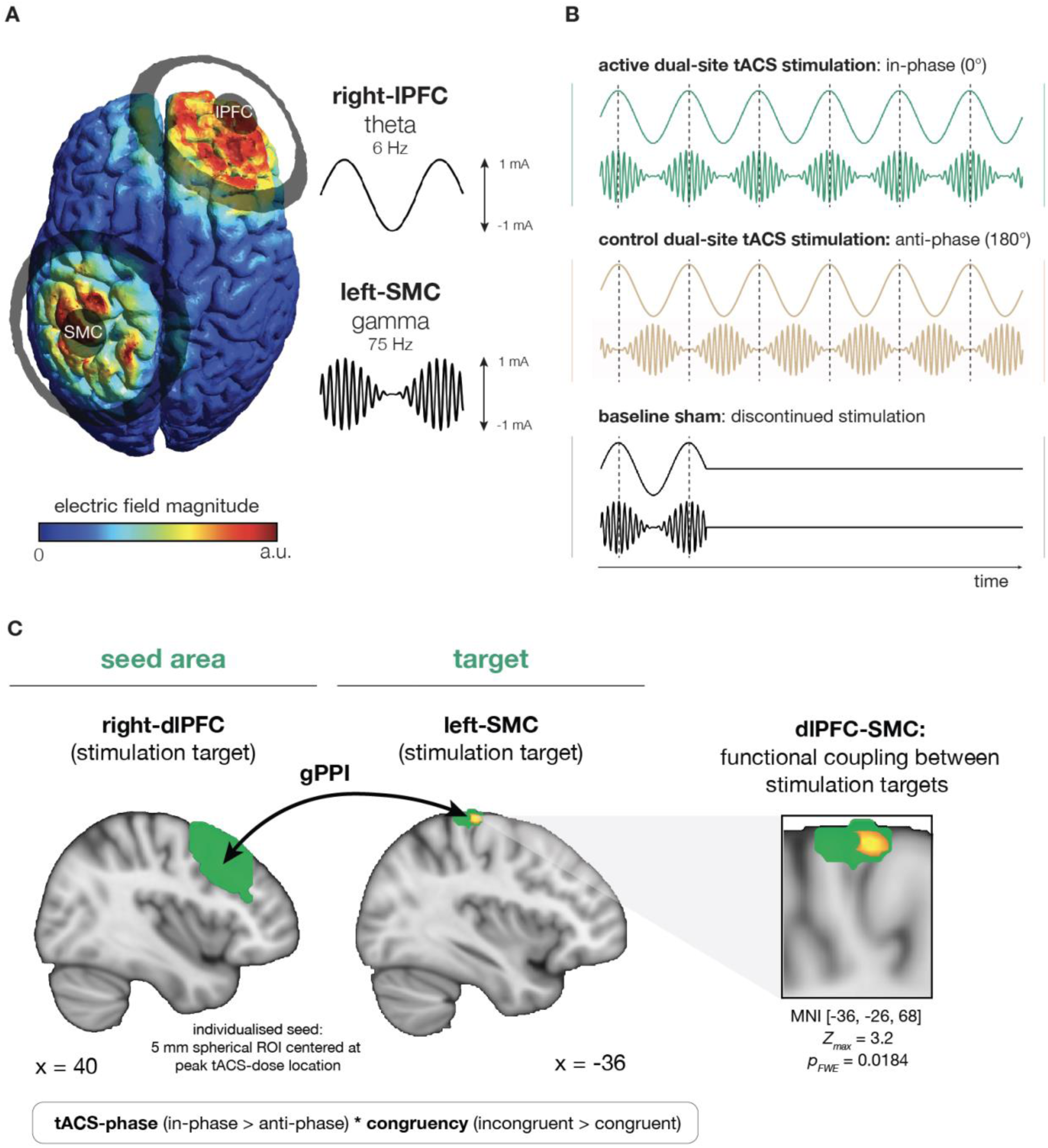
**Dual-site phase-amplitude coupled tACS modulates endogenous inter-regional coupling between stimulation targets. (A) Modeled current density distribution at cortical targets.** Two sets of concentric ring electrodes were placed over the right lPFC and left SMC. The current density model shows the spatial distribution of the electric field magnitude evoked by the high-definition montage at both cortical targets. **(B) Dual-site phase-locked tACS manipulations.** The stimulation conditions underwent a pseudo-random alternation throughout the experiment between active in-phase tACS (green), active control anti-phase tACS (brown), and baseline sham (black). Specifically, 75 Hz stimulation was administered over the SMC and its amplitude modulated by the phase of a 6 Hz stimulation over the lPFC. Modulation was either in-phase or anti-phase with the peaks of the 6 Hz lPFC stimulation. While both in-phase and anti-phase tACS may influence local target excitability, in-phase tACS is specifically designed to enhance endogenous phase-amplitude coupling between the dlPFC and SMC, a mechanism associated with successful emotional action control (Bramson et al., 2018, 2020a). The baseline sham condition involved a brief 10-second initial stimulation that was discontinued before the start of the first trial. **(C) tACS-phase effect on congruency-related dlPFC-SMC coupling.** In-phase vs. anti-phase tACS selectively increased congruency-related functional coupling between dlPFC (seed) and SMC (target) stimulation areas (in-phase > anti-phase * incongruent > congruent), confirming the effectiveness of the tACS-phase manipulation in modulating inter-regional functional connectivity supporting emotional action control.

### MRI

All participants underwent magnetic resonance imaging using a 3T MAGNETRON Prisma MR scanner (Siemens AG, Healthcare Sector, Erlangen, Germany), which has a 64-channel head coil with a top opening that allows the tACS electrode wires to be routed out of the back of the scanner bore. The scans obtained during the MR sessions were aligned with a standard brain atlas to guarantee a uniform field of view (FoV) throughout the days. Each scanning day involved taking around 1800 functional images using a multi-band six sequence, 2 mm isotropic voxel size, TR/TE = 1000 / 34 ms, flip angle = 60°, phase angle P >> A, including ten volumes with reversed phase encoding direction (A >> P) used to correct image distortions.

High-resolution structural images were acquired with a single-shot MPRAGE sequence using a GRAPPA acceleration factor of 2, TR/TE = 2400/2.13 ms, an effective voxel size of 1 mm isotropic, 176 sagittal slices, a distance factor of 50%, a flip angle of 8°, orientation A >> P, and a FoV of 256 mm.

### Behavior analyses

To examine effects of congruency (congruent, incongruent) and stimulation (in-phase, anti-phase, sham) on behavioral response accuracy (correct, incorrect), we employed mixed-effects logistic regression using the *lme4* package in R (Bates et al., 2015). The models accounted for inter-individual variability by incorporating random slopes, intercepts, and correlations across participants to adhere to the maximal random effects structure. Outputs of these models are log odds (‘*b*’). Effect sizes were reported as odds ratios (ORs) with 95% confidence intervals (CIs). Statistical significance for all analyses were evaluated at the ɑ < 0.05 criterion.

Mixed-effects logistic regression models included behavioral accuracy (correct, incorrect) as dependent variable, with the relevant experimental conditions (congruency, stimulation) and their interactions as predictors. The tACS-dose (see below; continuous variable; *Z*-scored) metric was added as covariate to the interaction.

We hypothesized that the congruency effect in error rates would decrease for in-phase tACS and that the size of the effect per participant would depend on the BOLD effect of tACS *vs*. sham, a measure of dose dependence that is orthogonal to the contrast of interest (in-phase *vs*. anti-phase). These expectations were preregistered at the Open Science Framework: (https://osf.io/j9s2z/).

### ROI masks

The sensorimotor cortex (SMC) mask was created as a 10 mm spherical ROI centered at MNI coordinates [-36, -30, 72], based on model-based estimates of current density distribution under the SMC electrode location, along with the anatomical location used for neuronavigation. The dorsolateral prefrontal cortex (dlPFC) mask was derived from the right middle frontal gyrus mask in the probabilistic Harvard-Oxford Atlas (included in FSL). The bilateral amygdala mask was created using the right and left amygdala masks from the Harvard-Oxford Atlas, thresholded at 25%.

### fMRI analyses – preprocessing

All image processing was performed using MELODIC 3.00, as implemented in FSL 6.0.0 (https://fsl.fmrib.ox.ac.uk; (Jenkinson et al., 2012)). Motion correction was applied using MCFLIRT (Jenkinson et al., 2002), while magnetic field distortions were corrected with TOPUP (Andersson et al., 2003). Functional images were rigid-body registered to the brain-extracted structural image using FLIRT (Jenkinson et al., 2002). Nonlinear registration to MNI 2 mm standard space was performed with FNIRT. Images were then spatially smoothed with a 5 mm Gaussian kernel, high-pass filtered at 100 s, and pre-whitened. Independent component analysis (ICA) was conducted with a maximum of 100 components (Beckmann and Smith, 2004), which were manually inspected to remove potential noise sources (Griffanti et al., 2017).

### fMRI analyses – GLM

Emotional action control effects on whole-brain BOLD activation were estimated by contrasting incongruent trials (approach angry and avoid happy) with congruent trials (avoid angry and approach happy) across all stimulation conditions (in-phase, anti-phase, and sham).

tACS-dose effects on the whole-brain BOLD signal were assessed by comparing stimulation conditions (in-phase and anti-phase) to sham across both congruency conditions. Individual estimates of prefrontal tACS-dose were extracted from the 98% peak-activated voxel within the Harvard-Oxford middle frontal gyrus (dlPFC) mask. This tACS-dose metric (active tACS > sham tACS) is orthogonal to both the tACS-phase manipulation and the emotional action control contrast. Because concurrent fMRI-EEG studies have shown that low-frequency brain rhythms (e.g. theta) are associated with reduced BOLD signal (Scheeringa et al., 2009; Algermissen et al., 2022), in line with their predominantly inhibitory role (Yu et al., 2023; Leung and Yim, 2025), we consider a net reduction in prefrontal BOLD signal as a marker of stronger tACS-dose (see also Bramson et al., 2020a). Additionally, an exploratory whole-brain correlation analysis was conducted to examine the relationship between individual variation in STAI trait anxiety (*Z*-scored) and the tACS-dose contrast.

tACS-phase effects on congruency-related changes in dlPFC coupling were examined using generalized psychophysiological interaction (gPPI) analysis (O’Reilly et al., 2012). We created an interaction contrast between congruency (incongruent > congruent), tACS-phase (in-phase > anti-phase), and the mean dlPFC BOLD time series. The mean dlPFC signal for the seed regressor was extracted from a spherical ROI with a 5 mm radius, centered at each individual’s peak tACS-dose location. tACS-phase effects on temporal BOLD signal coupling between dlPFC (seed) and SMC (target) were assessed using small volume correction. Cluster-based inference was performed using Gaussian Random Field Theory correction, as implemented in FSL’s cluster tool (Smith and Nichols, 2009). A cluster-defining threshold of |*Z*| = 2.3 and a corrected cluster significance threshold of *p* < 0.05 were applied to control for multiple comparisons within the ROI.

First- and second-level GLM analyses were performed in FSL 6.0.5 using FEAT (Jenkinson et al., 2012). At the first level, the model included 12 task regressors: approach angry, approach happy, avoid angry, and avoid happy, modeled separately for each tACS condition (in-phase, anti-phase, and sham). Each regressor covered the time interval from stimulus presentation until joystick movement onset. The model also included nuisance regressors for estimated head motion (six translation and rotation parameters), their temporal derivatives, and global signal time series from white matter and cerebrospinal fluid. First-level models from two separate stimulation sessions were combined using fixed effects analysis in FEAT (Jenkinson et al., 2012).

At the group level, effects were assessed using FLAME 1 with outlier de-weighting (Woolrich, 2008). Family-wise error correction was applied using cluster-based inference with a cluster-forming threshold of |*Z*| > 2.3, which provides a false error rate of approximately 5% when using FLAME 1 (Eklund et al., 2016).

## RESULTS

### Behavioral cost of emotional action control

Forty-nine participants, selected on high self-reported daily-life social anxiety (**Fig. 1A**), were instructed to use a joystick to approach happy and avoid angry faces (affect-congruent condition) or avoid happy and approach angry faces (affect-incongruent condition), **Fig. 1B**. The affect-incongruent condition requires participants to override automatic emotional action tendencies to avoid angry and approach happy faces (Roelofs et al., 2009a; Phaf et al., 2014). Expected congruency effects were observed in the baseline sham-tACS condition, with higher error rates for incongruent as compared to congruent responses (*b*_congruency_ = 0.35, 95% *Confidence Interval* (*CI*) [0.09, 0.62], *χ²*_(1)_ = 7.06, *p* = 0.0079) (Volman et al., 2011; Bramson et al., 2018, 2020a; Kaldewaij et al., 2021), **Fig. 1C**. Specifically, the odds of participants making an incorrect response were 42% higher in the incongruent as compared to congruent condition (*Odds Ratio* (*OR*) = 1.42, 95% *CI* [1.10, 1.85]). This shows that exerting emotional action control is behaviorally costly.

### Socially anxious individuals recruit dlPFC for emotional action control

The patterns of whole-brain neural activation largely aligned with previous findings using the same task in non-anxious individuals, showing increased activation during incongruent *vs*. congruent trials in bilateral precuneus [14, -70, 36], lingual gyrus [16, -74, -4], and right dorsal anterior cingulate cortex [8, 32, 28], as well as decreased activity in bilateral medial [-8, 34, -14] and lateral orbitofrontal cortex [-40, 28, -14; 24, 28, -14], temporal pole [-42, 16, -36], fusiform cortex [40, - 42, -18], amygdala [38, -4, -20], extending into hippocampus and parahippocampal gyrus (Volman et al., 2011; Bramson et al., 2018, 2020a), **Table 1** and **Fig. 1D**. However, highly anxious individuals exhibited more robust activation for incongruent trials in the dorsolateral prefrontal cortex (dlPFC; Brodmann areas 9/46d) [30, 36, 38], rather than in the lateral frontal pole (FPl) region previously found in non-anxious individuals (Bramson et al., 2020a; Kaldewaij et al., 2021; Lapate et al., 2022). The differences in emotional action control circuits between high-anxious and non-anxious individuals have been detailed in a separate paper (Bramson et al., 2023). These findings indicate that high-anxious individuals likely implement control using dlPFC rather than FPl as non-anxious individuals (Bramson et al., 2023).

**Table 1.**
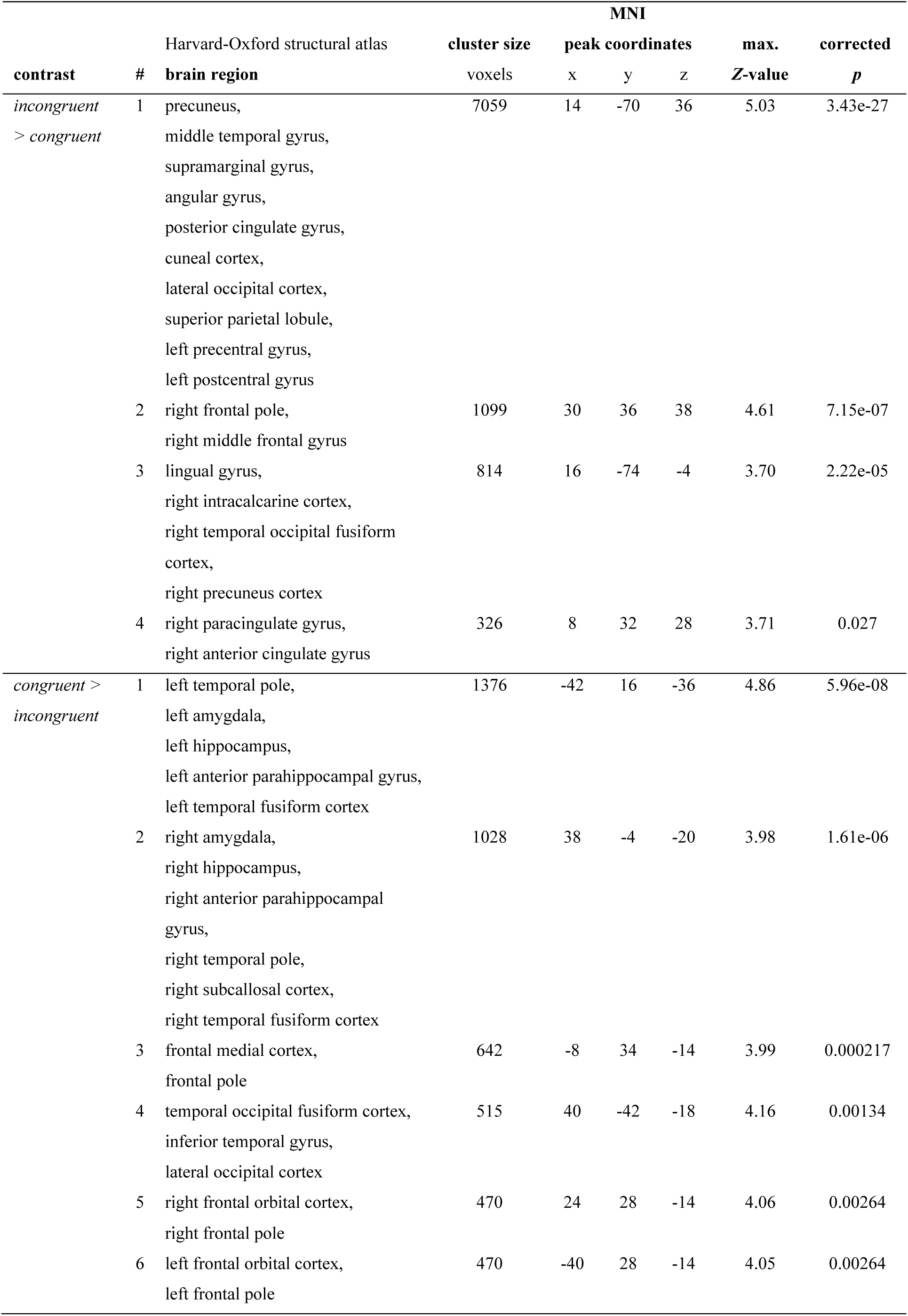
Group-level effects in the whole-brain BOLD-GLM analysis were assessed using FSL’s FLAME 1 mixed-effects model with automatic outlier de-weighting. Cluster-level inferences were family-wise error-corrected, applying a cluster-forming threshold of |Z| > 2.3 and p < 0.05.

### In-phase tACS strengthens dlPFC-SMC functional coupling

We used simultaneous tACS and fMRI to verify neural consequences of the tACS-phase manipulation, **Fig. 2A**. In-phase *vs.* anti-phase tACS (**Fig. 2B**) selectively enhanced functional connectivity between dlPFC and SMC target regions under the congruency challenge, as quantified using generalized psychophysiological interaction (gPPI) analysis (SMC cluster: *Z*_max_ = 3.2, *p*_fwe_ = 0.0184, MNI[-36, -26, 68]), **Fig. 2C**. This finding underscores the specificity of the in-phase tACS manipulation in strengthening inter-regional dlPFC-SMC coupling.

### Facilitation of emotional action control scales with dlPFC target engagement

Concurrent tACS-fMRI was used to study the underlying mechanisms of dual-site tACS effects, as well as to explain inter-individual variance in facilitation of behavioral performance depending on prefrontal stimulation reactivity. Following our pre-registered analysis pipeline (https://osf.io/j9s2z), we evaluated the cerebral effects of prefrontal tACS by examining BOLD activity in the region involved in implementing emotional action control in these high-anxious participants. Given that high-anxious individuals recruit dlPFC to solve emotional action control (Bramson et al., 2023), and that the dual-site tACS montage of this study evokes electrical fields covering dlPFC, we choose dlPFC BOLD activity as a metric of tACS dose-response. As in (Bramson et al., 2020a), the tACS-dose metric is obtained from a BOLD contrast (active tACS > sham tACS) that is orthogonal to both the tACS-phase manipulation and the emotional action control contrast. and we consider a net reduction in prefrontal BOLD signal as a marker of stronger tACS-dose (Scheeringa et al., 2009; Bramson et al., 2020a). Critically, we observe tACS-dose and tACS-phase specific enhancement of emotional action control (*b*_dlPFC-dose_ = -0.34, 95% CI [-0.66, - 0.02], *χ²*_(1)_ = 4.43, *p* = 0.0353), **Fig. 3A**. Participants with a more robust inhibitory response to prefrontal tACS exhibited greater improvement in emotional action control when receiving in-phase as opposed to anti-phase tACS. Planned post-hoc analyses on congruency-related error rates confirm significant dose-dependent effects for the in-phase (Spearman’s *ρ*_(47)_ = 0.37, *p* = .0092) but not for the anti-phase condition (Spearman’s *ρ*_(47)_ = -0.09, *p* = .5194). The direction and magnitude of the dual-site in-phase tACS intervention effects align with our previous report (Bramson et al., 2020a), and were specifically linked to tACS-dose measured over dlPFC. Namely, when extracting tACS-dose from FPl, a region previously reported to be decoupled from emotional action control in high-anxious individuals (Bramson et al., 2023), no significant tACS-induced changes in emotional action control were observed (*b*_FPl-dose_ = -0.12, 95% CI [-0.46, 0.22], *χ²*_(1)_ = 0.49, *p* = 0.4844). Further exploration of brain-symptoms correlations revealed a positive association between higher trait anxiety levels and stronger tACS-dose in dlPFC, **Fig. 3B**. In summary, the findings demonstrate that dual-site tACS can effectively enhance emotional action control in high-anxious individuals by targeting phase-amplitude coupling between dlPFC and SMC.

**Figure 3.**
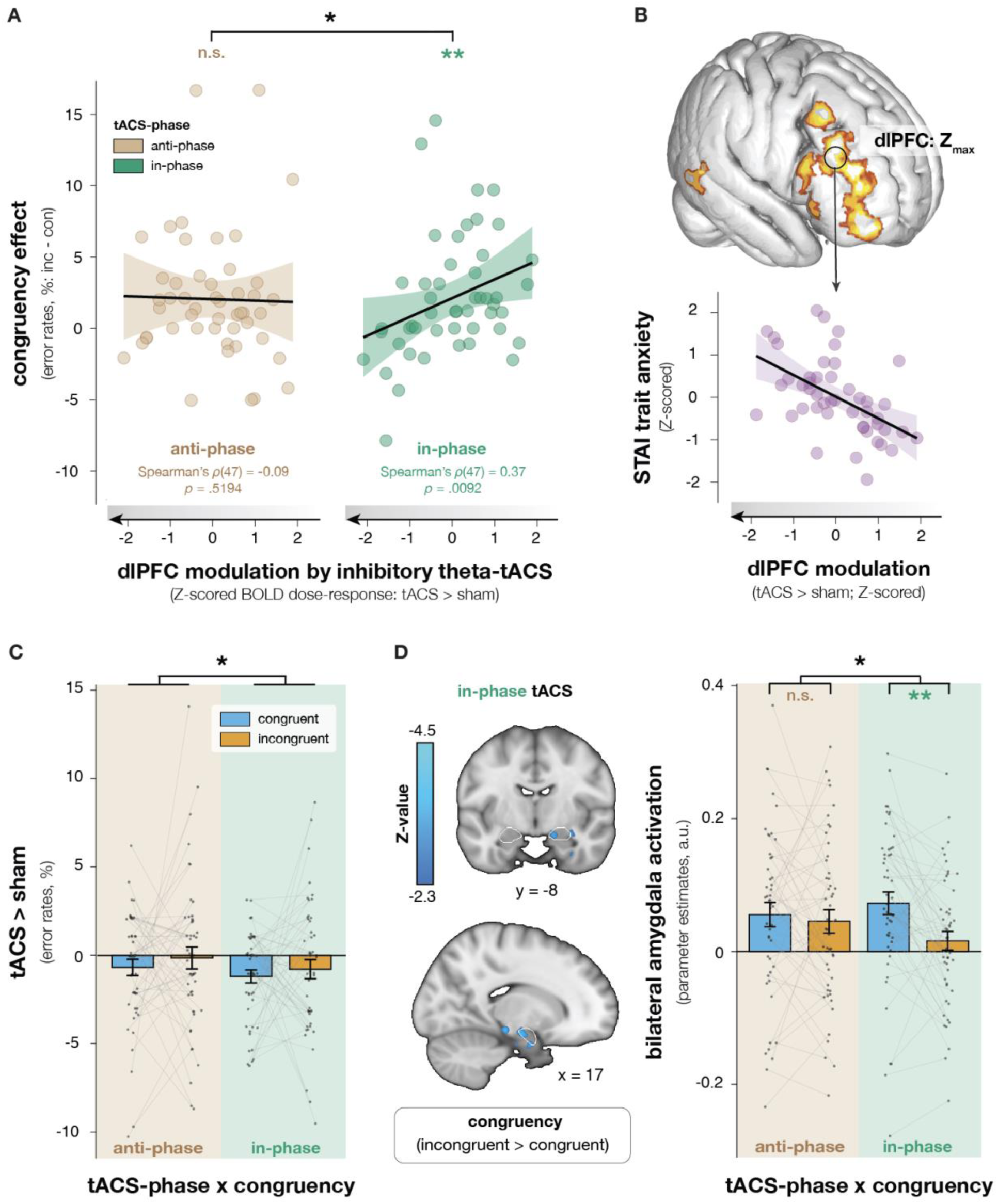
**Phase- and dose-specific effects of dual-site tACS on emotional action control in social anxiety. (A) In-phase tACS facilitation of emotional action control is dose-dependent.** Participants who exhibited stronger inhibitory theta-band stimulation responses in dlPFC, as evidenced by a decrease in BOLD signal (Scheeringa et al., 2011; Bramson et al., 2020a), demonstrated improved control over emotional actions (i.e., a reduced congruency effect) during in-phase tACS (green) compared to anti-phase tACS (brown). Black lines represent the fitted linear trend, with the shaded area indicating the associated 95% confidence interval. **(B) tACS-dose scales with trait anxiety.** Concurrent tACS-fMRI BOLD signal changes across the in-phase and anti-phase tACS conditions (tACS-dose: tACS > sham) correlated with trait anxiety, indicating a stronger inhibitory dlPFC dose-response in individuals with higher trait anxiety levels. To illustrate this relationship, we included a scatterplot depicting the association between trait anxiety scores and tACS-dose values extracted from the maximum Z-value location within dlPFC. **(C) In-phase tACS improves general behavioral performance independent of tACS-dose.** In-phase (green panel) vs. anti-phase tACS (brown panel) selectively enhanced response accuracy across congruent (blue) and incongruent (orange) conditions compared to sham. Bar plots represent mean and standard error of the mean for each condition, with individual participant means indicated by grey dots. **(D) In-phase tACS of dlPFC-SMC coupling modulates amygdala contribution to valence-based action selection.** Left: In the in-phase tACS condition, the right amygdala exhibited negative congruency-related BOLD signal (incongruent > congruent; cluster-level inferences corrected for multiple comparisons). White outlines indicate the anatomical bilateral amygdala mask used for extracting beta parameter estimates. Right: In-phase tACS (green panel) increased congruency-related amygdala modulation compared to anti-phase tACS (brown panel).

### In-phase tACS improves valence-based action selection

We also examined tACS-phase effects independent of tACS-dose. In-phase tACS, but not anti-phase tACS, results in a general behavioral performance enhancement compared to sham (*b*_tACS-phase_ = -0.16, *SE* = 0.08, *p* = 0.0403), **Fig. 3C**. Specifically, in-phase tACS reduces the odds of participants making an incorrect response by 22% across both congruency conditions compared to sham (*b*_in-phase_ = -0.25, 95% *CI* = [-0.39, -0.11], *p* = 0.0004; *OR* = 0.78, 95% *CI* = [0.67, 0.89]). In contrast, anti-phase tACS had no significant effect on behavioral accuracy (*b*_anti-phase_ = -0.10, 95% *CI* = [-0.25, 0.06], *p* = 0.2225). Consistent with our previous work (Bramson et al., 2020a), when individual differences in neural responsiveness to tACS were not controlled for, tACS-phase did not show a congruency-specific effect on performance (*χ^2^* = 0.85, *p* = 0.6548).

Next, we explored the neural mechanisms underlying the general improvement in valence-based action selection during in-phase tACS. Given prior evidence that emotion control involves amygdala dampening (Volman et al., 2011, 2013, 2016) and that suppression of task-irrelevant valence signals reduces amygdala engagement (Anticevic et al., 2010; Kanske et al., 2011; Cohen et al., 2016) we focused on amygdala activation (bilateral). We find a significant effect of tACS-phase on congruency-dependent amygdala modulation (*b*_congruency:tACS-phase_ = 0.05, 95% CI = [0.00, 0.09], *F*_(1, 96)_ = 4.62, *p* = 0.0341, *ηₚ²* = 0.05), **Fig. 3D**. Specifically, in-phase tACS resulted in stronger congruency-related modulation of amygdala activation (*b*_congruency_ = 0.06, *t*_(95)_ = 3.47, *p* = 0.0008) compared to anti-phase tACS, which showed no significant effect (*b*_congruency_ = 0.01, *t*_(95)_ = 0.62, *p* = 0.5355). Together, these findings suggest that in-phase tACS facilitation of dlPFC-SMC coupling enhances valence signal processing based on the affect-congruency of action goals, in a dose-response manner, and supports amygdala contribution to affect-congruent responses.

## DISCUSSION

This study demonstrates that in-phase dual-site tACS, targeted at facilitating endogenous theta-gamma phase-amplitude coupling between lPFC and SMC, enhances control over emotional action tendencies in highly socially anxious individuals. The main finding is that dual-site tACS effects, previously reported to improve control over automatic approach-avoidance tendencies in non-anxious participants (Bramson et al., 2020a), generalize to a clinically relevant population with altered control over daily-life social-emotional behavior (LSAS ≥ 30) (Mennin et al., 2002; Roelofs et al., 2009b; van Peer et al., 2009). Furthermore, this study shows that the dual-site tACS intervention in high-anxious males and females evokes effects in a different prefrontal territory (dlPFC) than in non-anxious males (FPl) (Bramson et al., 2020a). This observation fits the recently reported shift in the prefrontal circuit implementing emotional action control from FPl (non-anxious participants) to dlPFC (high-anxious participants) (Bramson et al., 2023). The current study demonstrates the possibility of improving emotional action control using dual-site tACS in high-anxious individuals via the facilitation of endogenous long-range phase-amplitude coupling. This finding extends our mechanistic understanding of approach-avoidance control in social anxiety and introduces rhythmic synchronization between dlPFC and SMC as a novel therapeutic tool.

### dlPFC-SMC phase-amplitude coupling supports emotional action control in social anxiety

This dual-site tACS intervention builds on prior studies that have demonstrated the significance of endogenous lPFC-SMC theta-gamma coupling in controlling automatic emotional action tendencies and in facilitating the implementation of alternative goal-directed actions (Bramson et al., 2018, 2020a). In this study, in-phase dual-site tACS could enhance interregional synchronization of neuronal excitability periods, biasing action selection in SMC according to valence-incongruent action goals in dlPFC (Voytek et al., 2015; Bramson et al., 2018; Weber et al., 2024). In one important detail, this putative mechanism differs from what has been observed in healthy individuals, where prefrontal control emerged from FPl (Bramson et al., 2020a) rather than dlPFC (this study). This difference is important for understanding the scope and the mechanism of the dual-site tACS intervention in social anxiety. Namely, FPl is a region that can flexibly integrate valence signals with approach-avoid action selection, allowing FPl to orchestrate goal-directed control over automatic emotional action tendencies (Bramson et al., 2020b; Lapate et al., 2022). However, FPl is overexcitable in high-anxious individuals (Bramson et al., 2023), and the evidence suggests dlPFC compensates for FPl when control over emotional actions is required (Bramson et al., 2023). In contrast with integrated valence-action goal representations observed in FPl, action goal representations in dlPFC are not modulated by valence (Lapate et al., 2022), consistent with relatively sparse mono-synaptic connections from amygdala to dlPFC (Petrides and Pandya, 1999; Folloni et al., 2019). In fact, dlPFC engagement could dampen amygdala processing via strong indirect projections to the ventromedial subgenual cingulate area 25 (Joyce et al., 2020, 2023), thereby providing a candidate mechanism for gating the contribution of emotional afferences to cognitive control (Berboth and Morawetz, 2021). Thus, the dlPFC can implement rule-based actions with minimal interference from emotional valence signals even in high anxiety, when emotional signals would saturate an already overexcitable FPl (Bramson et al., 2023). However, this may come at a cost: dlPFC-based emotional action control might be limited when emotional challenges co-occur with demands for cognitive control, e.g. during real-life situations. Accordingly, high-anxious individuals show reduced dlPFC efficiency during cognitive control compared to non-anxious individuals (Eysenck et al., 2022). It remains to be seen whether dual-site tACS in socially-anxious individuals can enhance dlPFC coordination with SMC and remain behaviorally effective even in the presence of multiple cognitive control demands.

### tACS BOLD dose-response in dlPFC scales with trait anxiety

The current study shows that the application of in-phase dual-site tACS results in behavioral effects proportional to BOLD signal evoked over dlPFC across tACS conditions, i.e., a dose-response effect. We also observe that individuals with higher trait anxiety exhibit more robust dlPFC responses to the tACS intervention. This finding corresponds to the observation that trait anxiety is associated with an anatomical shift of prefrontal emotional action control, from FPl to dlPFC (Bramson et al., 2023). The link between tACS response and trait anxiety signals that sub-optimal endogenous synchronization could underlie reduced emotion control performance in anxiety. Namely, the most robust effects of tACS are expected when weak endogenous oscillations can be facilitated (Krause et al., 2022), and in-phase dual-site tACS is expected to improve long-range communication by reducing noise within hypo-synchronized circuits (Wischnewski et al., 2023). Specifically, we suggest that trait anxiety might index individual levels of neural noise within the dlPFC-SMC circuit during emotional action control, a prediction that could be tested by measuring electrophysiological activity during emotional action control in high-anxious individuals (Bramson et al., 2018). This knowledge would be instrumental in facilitating the translation of dual-site tACS to clinical protocols, as it would enable using trait anxiety rather than concurrent tACS-fMRI to estimate individual dose-responses.

### Interpretational issues and future directions

It could be argued that the current findings are limited by the lack of electrophysiological observations of the consequences of tACS. In fact, the manipulation of dual-site tACS phase angle (in-phase *versus* anti-phase) enables us to relate the tACS effects to phase-specific changes in long-range phase-amplitude coupling, and exclude general changes in lPFC or SMC excitability. The phase manipulation provides the additional benefit of fully matching peripheral effects between in-phase and anti-phase stimulation, thereby ruling out alternative explanations related to transcutaneous entrainment or retinal stimulation (Schutter, 2016; Asamoah et al., 2019).

In this study, phase-specific dual-site tACS improvements in emotional action control are dose-related, an indication of high inter-individual variation in the effects of dual-site tACS on emotional action control. This variation makes the tACS effects physiologically plausible, since tACS effects depends on the electric field induced in the brain (Krause et al., 2019; Johnson et al., 2020) and on the interaction with ongoing endogenous oscillations (Krause et al., 2022). Future studies could use individualized current flow models to optimize stimulation protocols and enhance targeting accuracy to achieve more consistent tACS effects (Grover et al., 2023). However, these models do not capture state-dependent effects in the neural response to stimulation (Bradley et al., 2022), which could be particularly relevant for translational efforts in which the neural mechanisms underlying the disorder are not fully understood. Inter-individual variation in tACS efficacy can be exploited to identify factors that could maximize intervention response, in particular when used to enhance the effectiveness of existing treatments. This can include improving the ability to control emotional behavior during exposure therapy to confront threatening situations and temporarily interrupt the cycle of persistent avoidance.

## CONCLUSION

Building non-invasive neuromodulation interventions that facilitate cognitive function presents a major neuroscientific challenge due to the need to identify and enhance the underlying endogenous neural dynamics. Translating mechanistically informed neuromodulation interventions from healthy individuals to clinical populations poses a further challenge due to an incomplete understanding of the neural mechanisms underlying psychiatric disorders. Here, we successfully translate a mechanistically informed dual-site tACS intervention (Bramson et al., 2018, 2020a) to a clinically relevant high-anxiety population that shows impairments in daily-life social-emotional behavior. The findings highlight the advantage of combining cognitive neuromodulation interventions with neuroimaging during the early stages of clinical translation. The advantage of this multi-modal approach is that it offers a better understanding of the intervention mechanism, leading to a more precise characterization of inter-individual variation in intervention response. The results provide insights into the role of phase-amplitude coupling between lPFC and SMC in emotion control over social approach-avoidance behavior and contribute to developing targeted clinical interventions for anxiety.

## Acknowledgments

We thank Jesse Lam for his support with data collection.

## Data and materials availability

All behavioral and fMRI data, along with the analysis scripts, will be made available at https://doi.org/10.34973/jjjq-nr84.

